# High latitude, high tolerance: Northern juvenile Chinook salmon exhibit substantial thermal plasticity and acute heat tolerance

**DOI:** 10.64898/2026.11.04.764302

**Authors:** A. L. Wagner-Jones, S. A. Berkelhammer, L. Barreiros Scatamburlo, E. P. Williams, A. M. Snyder, M. J. H. Gilbert

## Abstract

In a rapidly warming North, Chinook salmon are facing rising river temperatures, increasing frequencies of heat waves, and greater annual thermal variation. These factors have putatively contributed to substantial declines in Alaska Chinook salmon abundance; understanding the thermal sensitivity and acclimation capacity of Chinook salmon at the northern extents of their range is therefore imperative. To this end, we acclimated juvenile Yukon River Chinook to five ecologically relevant summer temperatures (12.5, 15, 17.5, 20, and 22.5°C) under two feeding treatments of 100% or 40% satiation. We then assessed whole organism heat tolerance at rest and while swimming, along with the thermal sensitivity of heart rate and cardiac electrical activity. Yukon River Chinook substantially increased whole organism heat tolerance at rest and under exercise with warm acclimation. Cardiac heat tolerance also increased, and fish showed marked thermal compensation of maximum heart rate, resetting it to lower levels with warm acclimation. Feed treatment had a limited effect outside of peak heart rate and the transition temperatures of Q_10_. Warm acclimation was also associated with a shortening of the PR and QRS intervals and a lengthening of the QT interval. Across all acclimation groups, the PR interval plateaued to an extent far greater than heart rate itself, suggesting that limitations to AV conduction may limit cardiac function at high acute temperatures. Altogether, high latitude juvenile Chinook display remarkable heat tolerance and thermal plasticity, which may be preserved in association with the large seasonal, weather dependent, and spatial thermal variation Yukon River Chinook face during freshwater rearing and along with their extensive seaward and return migrations. The acute heat tolerance of Yukon River Chinook, which is comparable to or outperforming lower latitude Chinook and other salmonids, emphasizes the need for population-specific work for informed conservation efforts.

## Introduction

Warming rivers and oceans are presenting varied thermal challenges for Pacific salmon (*Oncorhynchus* spp.) across their broad latitudinal range; such changes are particularly pronounced in the North. Climate warming has caused water temperatures to rise substantially within the last century and the frequency of extreme heat waves to increase (Stillman, 2019). Many salmon populations across Alaska, British Columbia, the Northwest United States and California have experienced marked declines that are, at least in part, linked to changing thermal regimes both in the freshwater and marine environments (DeFilippo et al., 2026; Mantua et al., 2010; Mathes et al., 2010; Martins et al., 2011; Oke et al., 2020; Rand et al., 2006; Schindler et al., 2013; Schoen et al., 2023; Wainwright & Weitkamp, 2013). Yukon River Chinook (*O. tshawytscha*) in particular have experienced steady population declines since the 1980s, with record low abundances in recent years (Schindler et al., 2013; Schoen et al., 2023). These declines have prompted several federal fishery disaster declarations (Dunleavy, 2025; USDC, 2022, 2023). Further, the Alaska Department of Fish and Game and Fisheries and Oceans Canada have issued a historic joint seven-year moratorium on Yukon River Chinook salmon fishing (ADF&G & DFO, 2024). While direct causes remain unresolved, high temperatures and thermal variability may reduce juvenile condition at marine entry and future survival (DeFilippo et al., 2026; Mayer et al., 2024). Yukon River temperatures historically range from ∼1-4°C in winter to 15-18°C in summer, with peaks above 21°C (Fig. 1; AKTEMP, 2026; Eiler et al., 2023). Because juveniles generally rear in the river for at least one year before migrating to sea, they may be particularly vulnerable to heat stress, which has been documented in both adult and juvenile Yukon River Chinook salmon at 21°C (Bowen et al., 2020; Von Biela et al., 2020, 2023).

**Figure 1:**
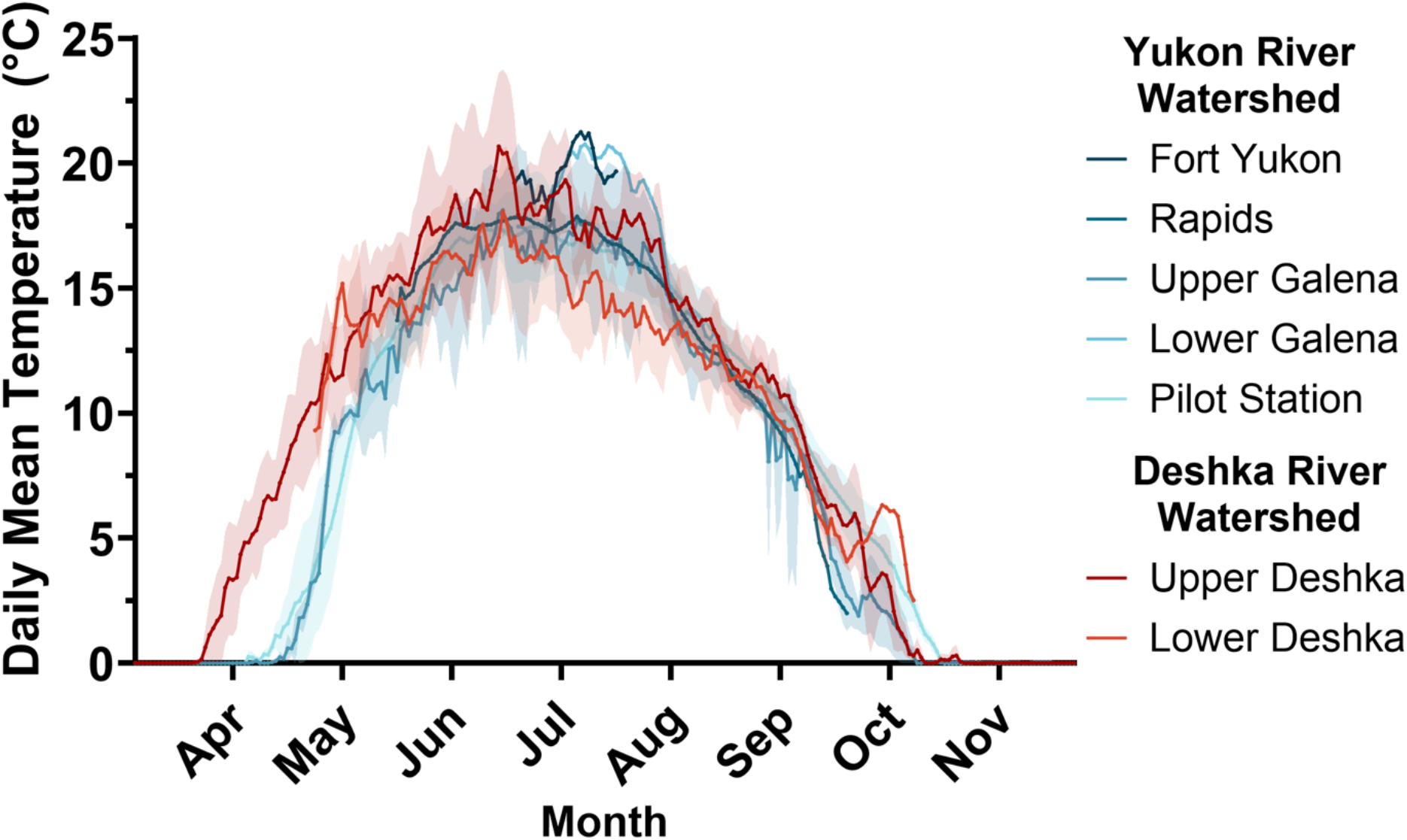
Historical mean daily temperatures for the Yukon River and Deshka River (± SD). Data were compiled for five stations along the Yukon River between 2002-2026 and two stations along the Deshka River between 2008-2019 from the Alaska Water Temperature Database (AKTEMP, 2026). **Alt text:** Graph depicting the mean daily water temperature from March to November of the Yukon River and Deshka River watersheds.

Heat tolerance and its plasticity vary across and within Pacific salmon species, with some populations being locally adapted to historical water temperatures and migratory challenges (Durhack et al., 2021; Eliason et al., 2011; Farrell, 2009; Van Wert et al., 2023; Zillig et al., 2023, 2025). Temperature sensitivity also varies across life stages, but in distinct ways among species and populations, with some showing the highest heat tolerance at the fry stage and others showing an increase after smoltification (Mayer et al., 2024; Metzger et al., 2026; Richter & Kolmes, 2005; Zillig et al., 2021). Food quality and availability can further modify thermal responses (Hardison et al., 2023; Hardison & Eliason, 2024). Such variation cautions against generalizing thermal limits among populations. Nonetheless, northern populations remain understudied despite rapid warming from Arctic amplification, resulting in a knowledge gap for evidence-based management.

In ectotherms including Chinook salmon, physiological functions generally improve with warming until functional and then critical upper thermal limits are reached (Farrell, 2016). Heart rate is one vital function that exemplifies this pattern in fish, rising with acute warming to a peak or plateau before becoming arrhythmic (Anttila, Couturier, et al., 2014; Gilbert et al., 2024). Because rising heart rate is central to increasing oxygen delivery during warming, limits to heart rate can constrain cardiorespiratory and whole organism performance at high temperatures (Farrell, 2009, 2016; Gilbert et al., 2024). Fish can undergo active thermal plasticity to adjust their physiology to better suit prevailing temperatures, broadening both cardiac and whole animal thermal performance (Gilbert et al., 2022; Hofmann & Todgham, 2010; Middleton et al., 2024; Sutcliffe et al., 2020). Whole animal and cardiac thermal plasticity have been widely documented among salmonids generally, as well as temperate and southern Chinook strains (e.g. Anttila et al., 2014; Eliason et al., 2011; Muñoz et al., 2015). However, it has not been examined in the northernmost Chinook salmon populations, which experience wide seasonal temperature ranges and marked acute heat events.

Thus, the overarching aim of this study was to characterize the acute thermal sensitivity and heat tolerance of juvenile Yukon River Chinook salmon. Further, we aimed to determine their potential for thermal plasticity over a range of environmentally realistic summer temperatures and the role of food availability in setting upper thermal limits. We hypothesized that, relative to more southerly distributed populations for which data exists, Yukon River Chinook salmon are either 1) adapted to colder average historical river temperatures and thus have reduced upper thermal limits and capacity for acclimation to warm temperatures (>18°C; Von Biela et al., 2020, 2023), or alternatively, 2) have a broad thermal performance, acclimation capacity and high heat tolerance associated with the impressive migratory challenges they face and the substantial seasonal and weather dependent thermal variation that occurs within the Yukon River basin (Eliason et al., 2011; Zillig et al., 2023). Furthermore, we hypothesized that reduced food availability, from excess to levels more common in nature, would decrease heat tolerance and the ability to acclimate high temperatures. To address these hypotheses, we reared juvenile Yukon River Chinook salmon at temperatures from 12.5 to 22.5°C and fed them to 100% or 40% satiation for >3 weeks. We then measured critical thermal maxima at rest (CTmax) and during swimming (CTswim) and the response of maximum heart rate to acute warming, and compared results with data for more southern strains.

## Methods

### Experimental animals & study design

This study was approved by the University of Alaska Fairbanks Institutional Animal Care and Use Committee (IACUC #2193686-3) and Alaska Department of Fish and Game (SF2025-127). The fish used in this study were wild origin juvenile Chinook salmon. Gametes were collected by the Alaska Department of Fish and Game from adults in the Salcha River within the Yukon River watershed and reared by the Ruth Burnett Sport Fish Hatchery (Fairbanks, Alaska) at ∼7°C for 7 months on commercial hatchery feed. For the present study, 402 juvenile Chinook salmon were obtained from the hatchery May 27^th^, 2025, and were housed in the Biological Research and Diagnostics Aquatics Facility at the University of Alaska Fairbanks (UAF). The fish were split evenly (∼20 fish per tank) across twenty 150L tanks, with four tanks per independent 750L recirculating system (5 systems total; each set to a particular acclimation temperature) in reverse-osmosis water remineralized with 50% Remineraliz-P (Brightwell Aquatics) and 50% Instant Ocean Sea Salt (Instant Ocean.; ∼300uS/cm, pH ∼7.0) After transfer, fish were held at 7°C for nine days on a 16:8 photoperiod and were transitioned to a chironomid larvae (bloodworm) diet before recording weight and length, and tagging fish on the dorsal and pectoral fins using Visible Implant Elastomer (Northwest Marine Technology). After tagging, systems were warmed by 2°C per day until the final treatment temperatures of 12.5, 15, 17.5, 20, and 22.5°C were achieved. The start of warming was staggered so that no more than two acclimation start dates occurred on the same day. All fish were fed a chironomid larvae (bloodworms) diet supplemented with mysis shrimp, better matching their natural diets, with two tanks of each acclimation temperature fed until satiation (food remaining in the bottom of the tank after 5 minutes) and the other two tanks receiving 40% of the ration mass consumed by the satiation-fed tanks normalized to the biomass in each tank. Weight and length measurements were collected at day 10, 20, and immediately following euthanization for use in an ongoing complementary bioenergetics study. Fish were fasted for 24 hours prior to any experiments and euthanization.

### Assessment of critical thermal maxima

Critical thermal maxima (CT_max_) was assessed on five randomly selected fish from each tank (n_total_ = 100; n = 20 per acclimation temperature). Fish were transferred to a 150L holding tank held at their acclimation temperature before warming began at a common rate of 18°C hr^-1^ (0.33°C min^-1^). This warming rate was selected to allow for direct comparison with juvenile Chinook and other Pacific salmon from lower latitudes for which data exists. CT_max_ was identified as the temperature at which the fish lost equilibrium. Fish were recovered at their acclimation temperature or at ∼10°C below their CT_max,_ whichever was higher, for 5-10 minutes before being returned to their holding tank. CT_swim_ was also assessed on five individuals at a time but only for the 100% satiation treatment due to time constraints (n_total_ = 50; n = 10 per acclimation temperature). To begin CT_swim_ assessments, fish were transferred to a 30L swim tunnel (Loligo Systems, Viborg, Denmark) at their acclimation temperature and allowed to habituate to the swim tunnel at speed of ∼1 body length s^-1^ for 30 minutes. Flow rate was then increased to ∼80% (50.1 cm s ^-1^; 5.2 body lengths s^-1^) of the critical sustained swimming speed (*U*_crit_) for fish in the 12.5°C acclimation group (estimated in complementary experiments) and warming began at 10°C hr^-1^ (0.17°C min^-1^) to match the heart rate methodology (see below). CT_swim_ was identified as the temperature at which the fish could not continue to swim. After reaching CT_swim_, fish were recovered as described for CT_max_.

### Assessment of cardiac thermal sensitivity

The response of maximum heart rate (*f*_Hmax_) to acute warming was assessed on six to ten fish per acclimation temperature and feed treatment (n_total_ = 86; Table 1) following an established protocol (Gilbert et al., 2024) and using methods similar to those used on juvenile Chinook salmon of more southern origin (Muñoz et al., 2015), salmonids (e.g. Adams et al., 2022; Anttila et al., 2013, 2014; Casselman et al., 2012; Chen et al., 2013, 2015; Gilbert et al., 2022; Gilbert & Farrell, 2021; Pettinau et al., 2022; Williams et al., 2025) and other fish species (Drost et al., 2014, 2016; Haverinen et al., 2018; Kraskura et al., 2023). Briefly, fish were anesthetized in buffered tricaine methanesulfonate (TMS; 100 mg L-1 induction, 70 mg L-1 maintenance with gill irrigation; Syndel, Ferndale, USA), fitted with subdermal ECG electrodes, and injected with atropine (1.2 mg kg-1) and isoproterenol (4 mg kg-1; Sigma-Aldrich, St. Louis, USA) to pharmacologically induce maximum heart rate (*f*_Hmax_; Gilbert et al., 2024). After 20 min, the water was warmed at a rate of 10°C hr^-1^ (0.17°C min^-1^) in a stepwise fashion of 1°C every 6 min. Fish were removed after the onset of cardiac arrhythmia, which was identified as missing beats. Fish were euthanized, and weight and length were recorded.

**Table 1:** Sample size (n), mass (g), fork length (mm), and Fulton’s condition factor (K) for juvenile Chinook salmon fed to 40 or 100% satiation and acclimated to 12.5, 15, 17.5, 20, and 22.5°C. Feeding treatments are combined unless specified. Significant differences between feeding treatments are denoted with asterisks (*). Data are presented as mean ± SE. Model statistics are presented in Table S1.

|  | **Acclimation Temperature (°C)** |  |  |  |  |
|---|---|---|---|---|---|
|  | **12.5°C** | **15°C** | **17.5°C** | **20°C** | **22.5°C** |
| **Sample Size** | 18 | 16 | 16 | 16 | 18 |
| **40%** | 10 | 8 | 8 | 6 | 10 |
| **100%** | 8 | 8 | 8 | 10 | 8 |
| **Mass (g)** | 12.3 ± 0.5 | 11.6 ± 0.5 | 11.8 ± 0.5 | 11.0 ± 0.6 | 11.8 ± 0.6 |
| **Length (mm)** | 101.4 ± 1.4 | 99.7 ± 1.7 | 100.1 ± 1.4 | 97.5 ± 2.2 | 100.5 ± 1.4 |
| **Condition (K)** | 1.17 ± 0.02 | 1.16 ± 0.02 | 1.17 ± 0.03 | 1.18 ± 0.03 | 1.17 ± 0.07 |
| **40%** | 1.12 ± 0.02* | 1.14 ± 0.03* | 1.14 ± 0.05* | 1.08 ± 0.02* | 1.11 ± 0.02* |
| **100%** | 1.23 ± 0.04 | 1.18 ± 0.04 | 1.2 ± 0.02 | 1.24 ± 0.03 | 1.24 ± 0.16 |

The *f*_Hmax_, the peak *f*_Hmax_ during acute warming (peak *f*_Hmax_), the temperature at peak *f*_Hmax_ (T_peak_), and temperature at the onset arrhythmia (T_arr_) were identified by analysis of the ECG recordings using LabChart (ADInstruments, Colorado Springs, CO, USA). The incremental thermal sensitivity quotient (Q_10_) for *f*_Hmax_ during warming was calculated for each individual over every 2°C increment (Equation 1: R = *f*_Hmax_ in beats min^-1^) to characterize the thermal sensitivity of heart rate. The temperature at which incremental Q_10_ dropped and stayed below 2.0 (T_Q10<2_) was identified as an early indicator of the decline in the ability to increase *f*_Hmax_ to support elevated oxygen transport during warming (Eliason and Anttila, 2017; Gilbert et al 2024).

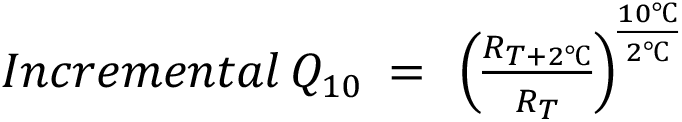

The acclimatory effects on key thermal tolerance metrics including CT_max_, CT_swim,_ T_peak_, and T_arr_ were calculated using Equation 2 where the difference in thermal tolerance between two acclimation temperatures (2 and 1) is divided by the difference in acclimation temperature.

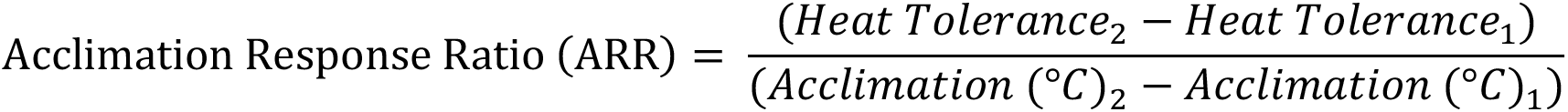

Detailed ECG waveform analysis was completed to assess electrical changes in the cardiac cycle between acclimation temperatures and with acute warming using the LabChart ECG module as described by Williams et al. (2025). Briefly, at the same 30s or ten beats where *f*_Hmax_ was taken, the P wave, QRS complex, and T wave were marked at their start, end, and peaks using automatic detection with manual adjustment as needed.

### Statistical analysis

Statistical analyses were completed in R using the lme4, glmmTMB, emmeans, and MuMIn packages (R Core Team, 2024; Bates et al., 2015; Brooks et al., 2017; Lenth & Piaskowski, 2026; Bartoń, 2026). Analysis of *f*_Hmax_ and incremental Q_10_ was performed using linear mixed effects models (LMER) and generalized linear mixed effects models (GLMER) with body mass as a covariate, fish ID as a random factor, and *f*_Hmax_ as a function of acclimation temperature, acute temperature, and/or feed treatment. For *f*_Hmax_ analysis, values were truncated at T_peak_ to characterize the increasing phase of the response to warming. The appropriate model was selected using the following pipeline: A global model was fitted with all interactions before all model permutations were generated using ‘dredge’. The top model was selected by AIC and if the difference in AIC between models was <2.0 the simplest model was chosen. Before modeling, assumptions (i.e. a normal data distribution) were tested using the Shapiro-Wilk test and analysis of diagnostic plots; if assumptions were met, a LMER was used and if not, a GLMER with gamma family and a log link was selected. Once the best model was selected an analysis of variance (ANOVA) output was generated followed by a Holm adjusted pairwise comparisons when significant effects were identified.

Visualizations were completed in GraphPad Prism v.8.3 (GraphPad Software, San Diego, USA). To better depict relative changes in the RR and PR intervals with warming, we normalized each to their initial values. We then plotted that normalized data with segmented linear regressions (SLR) to show the extent to which responses diverged at high acute temperatures with the assumption that the decline in RR and PR intervals should be similar with warming if PR interval was not limiting the frequency of contraction (Haverinen & Vornanen, 2020).

Two individuals were excluded from the entire dataset as statistical outliers (>1.5x outside the interquartile range) and because of early indications of poor sample preparations (highly variable QRS amplitude), but exclusion of these individuals did not meaningfully alter our conclusions (Figure S2). Two additional individuals are not represented in the detailed ECG dataset (RR, PR, QRS, and QT intervals; Figure 4 & 5) as waveforms outside of the QRS complex could not be clearly identified and fourteen individuals are not represented in the QT interval dataset as the T wave could not be confidently identified (e.g. low amplitude, noise).

## Results

### Fish size and condition

At the time of sampling, Chinook salmon had a mass of 11.7 ± 2.2 g and fork length of 99.9 ± 6.6 mm (Table 1). Including mass as a covariate did not improve *f*_Hmax_ models, and mass did not markedly differ between acclimation groups or feed treatments within acclimation temperatures (Acclimation: F_(4, 74)_ = 0.909, p = 0.4636, feed treatment: F_(1, 74)_ = 3.365, p = 0.071; Table 1, Table S1). However, as expected, Fulton’s condition factor was reduced in the 40% than 100% feed treatment (*χ*²(1) = 7.888, p = 0.005; Table 1, Table S1).

### Whole organism heat tolerance

Whole organism heat tolerance at rest (CT_max_) and under exercise (CT_swim_), increased with acclimation temperature (*χ*²(4) = 933.270, p < 0.001 and F_(4, 44)_ = 12.084, p < 0.001, respectively; Fig. 4, Table S2). Between 12.5 and 22.5°C, CT_max_ increased by 2.4°C (28.6 ± 0.3 to 31.0 ± 0.2 °C; *t*(93) = -27.130, p < 0.001) and differed between all acclimation temperatures but not between feed treatments (Fig. 4, Table S2). The acclimation response ratio (ARR) for CT_max_ between 12.5 and 22.5°C was 0.24 with ARR decreasing as acclimation temperature increased (ARR 12.5 *vs* 15°C: 0.34, 15 *vs* 17.5°C: 0.27, 17.5 *vs* 20°C: 0.21, 20 *vs* 22.5°C: 0.15). CT_swim_ increased more substantially, by 5.1°C between 12.5 and 22.5°C (23.2 ± 1.1 to 28.3 ± 1.1°C; *t*(44) = -5.579, p < 0.001) with CT_swim_ at 12.5°C being lower than that for all other acclimation temperatures (Fig. 4, Table S2). The ARR of CT_swim_ from 12.5 to 22.5°C was high at 0.5, but much of the increase occurred between acclimation temperatures of 12.5 to 17.5°C (ARR 12.5 *vs* 15°C: 1.34, 15 *vs* 17.5°C: 0.65, 17.5 *vs* 20°C: 0.01, 20 *vs* 22.5°C: 0.02).

### Cardiac heat tolerance and thermal sensitivity

As expected, all fish increased *f*_Hmax_ with acute warming up to a peak (peak *f*_Hmax_ at T_peak_) and variation in *f*_Hmax_ among individuals within an acclimation treatment was low (Fig. 2A and B; Fig. S3). In all but two fish the onset of cardiac arrhythmia (T_arr_) occurred within 1°C of T_peak_ with little decline in *f*_Hmax_ preceding T_arr_ (Fig. 2). Chinook salmon exhibited a robust resetting of *f*_Hmax_ with warm acclimation, resulting in lower *f*_Hmax_ at initial and intermediate acute exposure temperatures with an increase in acclimation temperatures from 12.5 and 22.5°C fish (e.g., - 15.4% at 12.5°C; t(76) = 10.518, p < 0.001; Fig. 2, Table 2). Warm acclimated fish also reached greater peak *f*_Hmax_ (6.4% increase; *χ*^2^(4) = 30.503, p < 0.001; Table 2, Table S2). The higher peak *f*_Hmax_ and reduced *f*_Hmax_ at low temperatures meant that the total increase in *f*_Hmax_ with acute warming (Δ*f*_Hmax_) increased by 32.0% with acclimation temperature from 12.5 to 22.5°C (*χ*^2^(4) = 9.470, p = 0.050; Table 2, Table S2).

**Figure 2:**
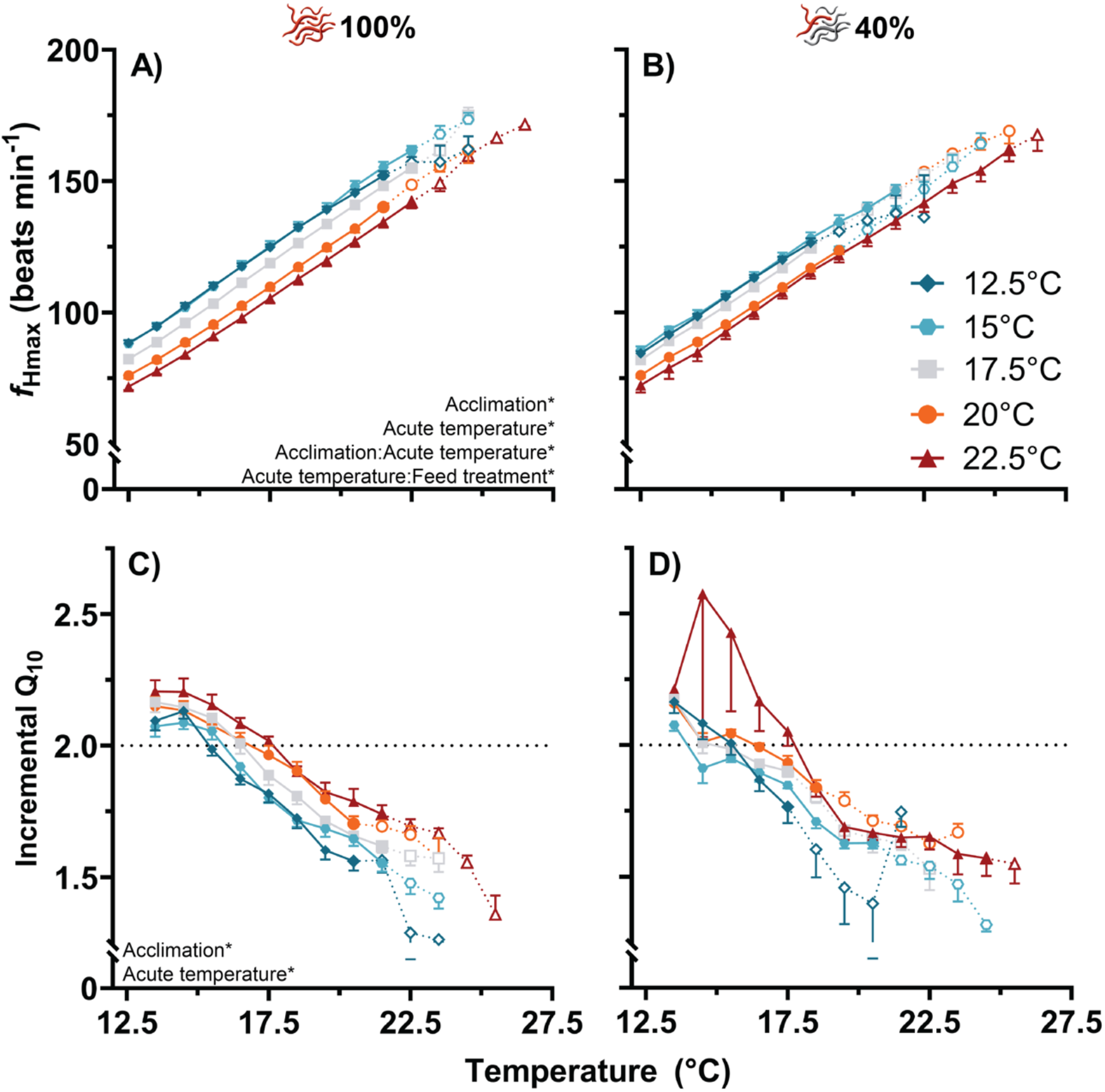
Response of maximum heart rate (*f*_Hmax_) and incremental Q_10_ to acute warming for juvenile Chinook salmon acclimated to 12.5, 15, 17.5, 20, and 22.5°C under two feeding treatments of 100% (A and C) or 40% (B and D) of satiation. Acute warming began at 12.5°C and continued at 10°C hr^-1^ until the onset of arrhythmia. Data are presented as mean ± SE. The dotted line and open symbols denote the point at which two individuals have been removed after the onset of arrhythmia. Significant fixed effects are noted on the graph, and further model statistics are presented in Table S3. **Alt text:** Graphs depicting the response of maximum heart rate and incremental Q_10_ to acute warming and the effect of acclimation, labeled A to D with annotations and statistical values.

**Table 2:** Heart rate (*f*_Hmax_; beats min^-1^) at 12.5, 17.5, and 22.5°C, peak *f*_Hmax_, and the change in *f*_Hmax_ from 12.5°C to peak *f*_Hmax_ (Δ*f*_Hmax_) for juvenile Chinook salmon fed to 40 or 100% satiation and acclimated to 12.5, 15, 17.5, 20, and 22.5°C. Acute warming began at 12.5°C and continued until the onset of arrythmia. Feeding treatments are combined unless specified. Dissimilar letters indicate significant differences between acclimation treatments and significant differences between feeding treatments are denoted with asterisks. Data are presented as mean ± SE. Model statistics are presented in Table S2.

|  | **Acclimation Temperature (°C)** |  |  |  |  |
|---|---|---|---|---|---|
|  | **12.5°C** | **15°C** | **17.5°C** | **20°C** | **22.5°C** |
| **12.5°C** | 86.2 ± 0.8<sup>a</sup> | 86.9 ± 0.9<sup>a</sup> | 82.1 ± 0.7<sup>b</sup> | 76.1 ± 0.7<sup>c</sup> | 72.2 ± 1.3<sup>d</sup> |
| **17.5°C** | 122.1 ± 1.0<sup>a</sup> | 123 ± 1.5<sup>a</sup> | 117.8 ± 0.9<sup>b</sup> | 109.7 ± 1.0<sup>c</sup> | 107 ± 1.5<sup>c</sup> |
| **22.5°C** | 152.1 ± 3.2<sup>abc</sup> | 158.7 ± 1.6<sup>a</sup> | 154 ± 1.1<sup>ab</sup> | 148.1 ± 1<sup>bc</sup> | 141.8 ± 1.9<sup>c</sup> |
| **40%** | 143.6 ± 6.5* | 153.5 ± 2.1* | 152.3 ± 1.4* | 146.9 ± 2.0* | 141.7 ± 3.1* |
| **100%** | 157.2 ± 1.6 | 161.7 ± 1.6 | 155.2 ± 1.6 | 148.6 ± 1.2 | 142 ± 2.3 |
| **Peak *f*<sub>Hmax</sub>** | 149.9 ± 2.7 | 161.6 ± 3.2 | 156.2 ± 3.5 | 150.8 ± 4.0 | 159.5 ± 4.3 |
| **40%** | 142.5 ± 3.3* | 157.7 ± 4.0* | 150.8 ± 5.2* | 147.2 ± 7.6* | 161.6 ± 6.5* |
| **100%** | 157.8 ± 2.0 | 165.6 ± 4.7 | 161.1 ± 4.1 | 152.7 ± 4.4 | 156.9 ± 5 |
| **Δ*f*<sub>Hmax</sub>** | 62.1 ± 2.7 | 72.7 ± 3.2 | 71.3 ± 3.8 | 72.0 ± 4.1 | 82.0 ± 5.5 |

Further, the slope of heart rate over acute temperature was significantly greater in the 100 than 40% feeding group, leading to generally higher heart rates and a greater Δ*f*_Hmax_ during acute warming (*χ*²(1) = 8.333, p = 0.004; Fig. 2B, Table 2, Table S2). Fish fed to 100% satiation reached significantly higher peak *f*_Hmax,_ than those at 40% satiation (+6.28%; *χ*^2^(1) = 4.848, p = 0.028; Table 2, Table S2). However, no pairwise comparisons of Δ*f*_Hmax_ or peak *f*_Hmax_ were significant between feed treatments at any single acclimation temperature (Table 2, Table S2).

As expected, incremental Q_10_ decreased during acute warming for all treatment groups. The greater increase in *f*_Hmax_ following warm acclimation was apparent in incremental Q_10_ values which were elevated over initial and intermediate temperatures (Fig. 2C and D; Q_10_ > 2.0; *χ*²(4) = 31.249, p < 0.001; Table S3). Between 12.5 and 22.5°C, T_Q10_ increased by 12.63% (+2.0°C; *t*(76) = -5.501, p < 0.001; Fig. 3, Table S2 and S5). Unlike *f*_Hmax_, we found weak evidence that food availability reduced incremental Q_10_ (p = 0.054; Table S3) during warming. Congruently, food-limitation did result in a lower T_Q10_ (F_(1, 76)_ = 4.660, p = 0.034; Fig. 2D, Fig. 3, Table S5) and appeared to increase variation between and within acclimation groups, especially at acute temperatures approaching their upper thermal limits.

**Figure 3:**
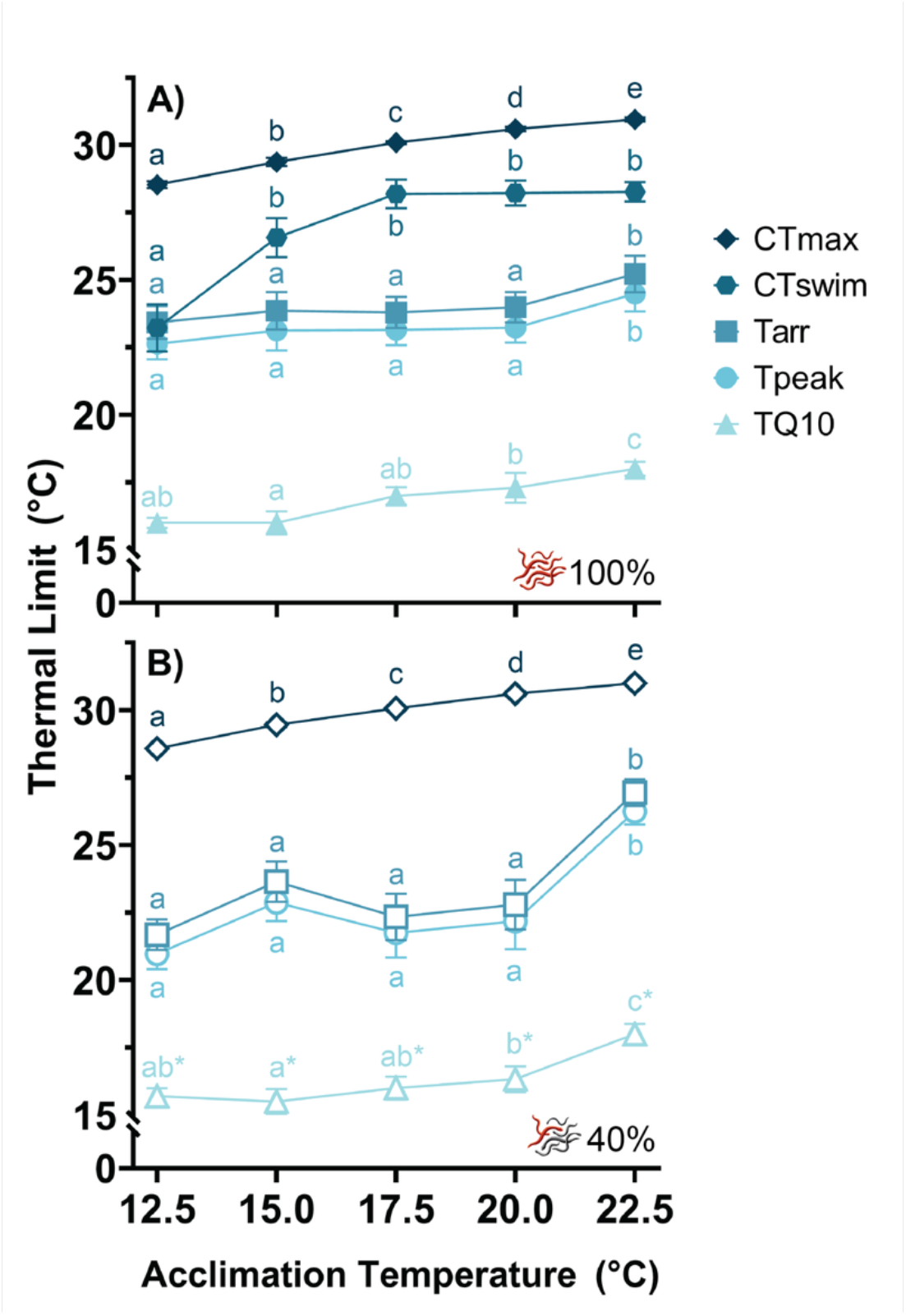
Effect of acclimation on transition and critical cardiac and whole animal temperatures for juvenile Chinook salmon acclimated to 12.5, 15, 17.5, 20, and 22.5°C and fed to 40 or 100% satiation. Transition and critical temperatures include critical thermal maxima static (CT_max_) and under exercise (CT_swim_), temperature at peak *f*_Hmax_ (T_peak_), temperature at arrhythmia (T_arr_) and temperature at which incremental Q_10_ dropped and remained below 2.0 (T_Q10_). Data are presented as mean ± SE with dissimilar letters indicating significant differences between acclimation temperatures and asterisks denoting significant differences between feed treatments. Model statistics are presented in Table S2. **Alt text:** Graphs depicting the increase of whole animal and cardiac thermal limits with acclimation temperature, with statistical values.

In addition to the increase in T_Q10_, warm acclimation was associated with an increase the upper thermal limits (T_peak_, T_arr_) of juvenile Chinook salmon (T_peak_: *χ*²(4) = 30.503, p < 0.001, T_arr_: F_(4, 76)_ = 8.036, p < 0.001; Fig. 3, Table S2 and S5). Between 12.5 and 22.5°C acclimation groups, T_peak_ increased by 12.36% (+2.7°C; *t*(76) = -5.216, p = 0.001) and T_arr_ by 11.82% (+2.7°C; *t*(76) = -5.313, p < 0.001; Fig. 3, Table S2 and S5). Correspondingly, T_peak_ and T_arr_ both displayed an ARR of 0.27 between 12.5 and 22.5°C, although the overarching acclimatory response was mainly driven by large increases at the extremes of the acclimation temperatures included in the study (T_peak_: ARR 12.5 *vs* 15°C: 0.52, 15 *vs* 17.5°C: -0.22, 17.5 *vs* 20°C: 0.16, 20 *vs* 22.5°C: 0.62; T_arr_: ARR 12.5 *vs* 15°C: 0.52, 15 *vs* 17.5°C: -0.28, 17.5 *vs* 20°C: 0.19, 20 *vs* 22.5°C: 0.63). As absolute values of T_peak_ and T_arr_ were closely linked, the thermal acclimatory responses were nearly identical. T_Q10_ also displayed variation of ARRs between acclimation groups, but to a far lesser degree and with a slightly differing trend with an overall ARR of 0.20 between 12.5 and 22.5°C (ARR 12.5 *vs* 15°C: -0.03, 15 *vs* 17.5°C: 0.30, 17.5 *vs* 20°C: 0.18, 20 *vs* 22.5°C: 0.36).

### Response of the cardiac cycle to acclimation

As *f*_Hmax_ rose during warming, the PR and QT intervals declined with the RR interval as expected, however, QRS duration increased slightly (Fig. 4). When compared at common low and intermediate temperatures, the duration of the RR interval increased progressively with acclimation temperature, corresponding with the changes in *f*_Hmax_ described above (Fig. 2A-B, Fig. 4). The QT interval duration did not vary with acclimation temperature, but QRS and PR intervals did (*χ*^2^(4) = 14.229, p = 0.007 and *χ*^2^(4) = 24.600, p < 0.001, respectively; Fig. 4, Table S4). Specifically, despite RR interval being longest in the 22.5°C acclimation group, the QRS and PR intervals were the shortest. Further, the slope of the PR interval against acute temperature significantly got progressively steeper with increasing acclimation temperature (*χ*^2^(4) = 24.600, p < 0.001; Fig. 4, Table S4) better aligning with corresponding declines in RR interval (Fig. 5). This difference was well captured by segmented linear regressions of normalized RR and PR intervals against acute temperatures. This analysis revealed that the PR interval plateaued to a greater extent than RR interval at temperatures much lower than T_peak_ for fish in all acclimation treatments (Fig. 5A-E) and that PR interval decreased to a greater extent in warm acclimated fish (Fig. 5F). No component of the ECG waveform varied with feed level so both treatments are combined for presentation (QRS: *χ*^2^(1) = 0.387, p = 0.534, PR: *χ*^2^(1) = 2.438, p = 0.118, QT: *χ*^2^(1) = 0.077, p = 0.782; Fig. 4, Table S4).

**Figure 4:**
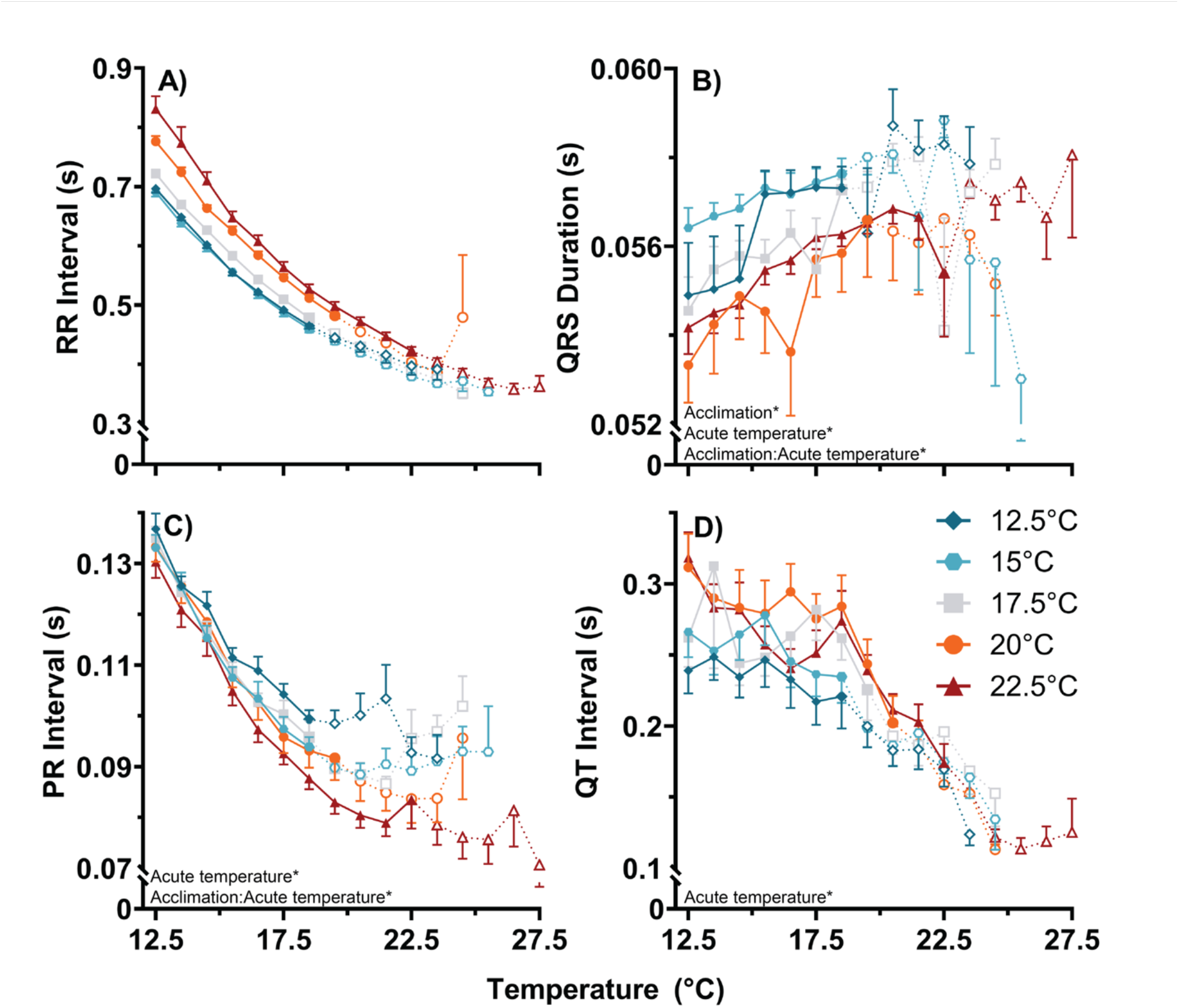
Effect of acclimation on components of the cardiac cycle under acute warming in maximally stimulated juvenile Yukon River Chinook salmon. Specifically, the A) RR, B) QRS, C) PR, and D) QT intervals were examined in Chinook salmon acclimated to 12.5, 15, 17.5, 20, and 22.5°C. Acute warming began at 12.5°C and continued at 10°C hr^-1^ until the onset of arrhythmia. Data are presented as mean ± SE and asterisks indicate significant model effects. The dotted line and open symbols denote the point at which two individuals have been removed after the onset of arrhythmia. Model statistics are presented in Table S4. **Alt text:** Graphs depicting the response of intervals of the cardiac cycle to acute warming and the effect of acclimation. Graphs are labeled A to D with detailed annotations and statistical values.

**Figure 5:**
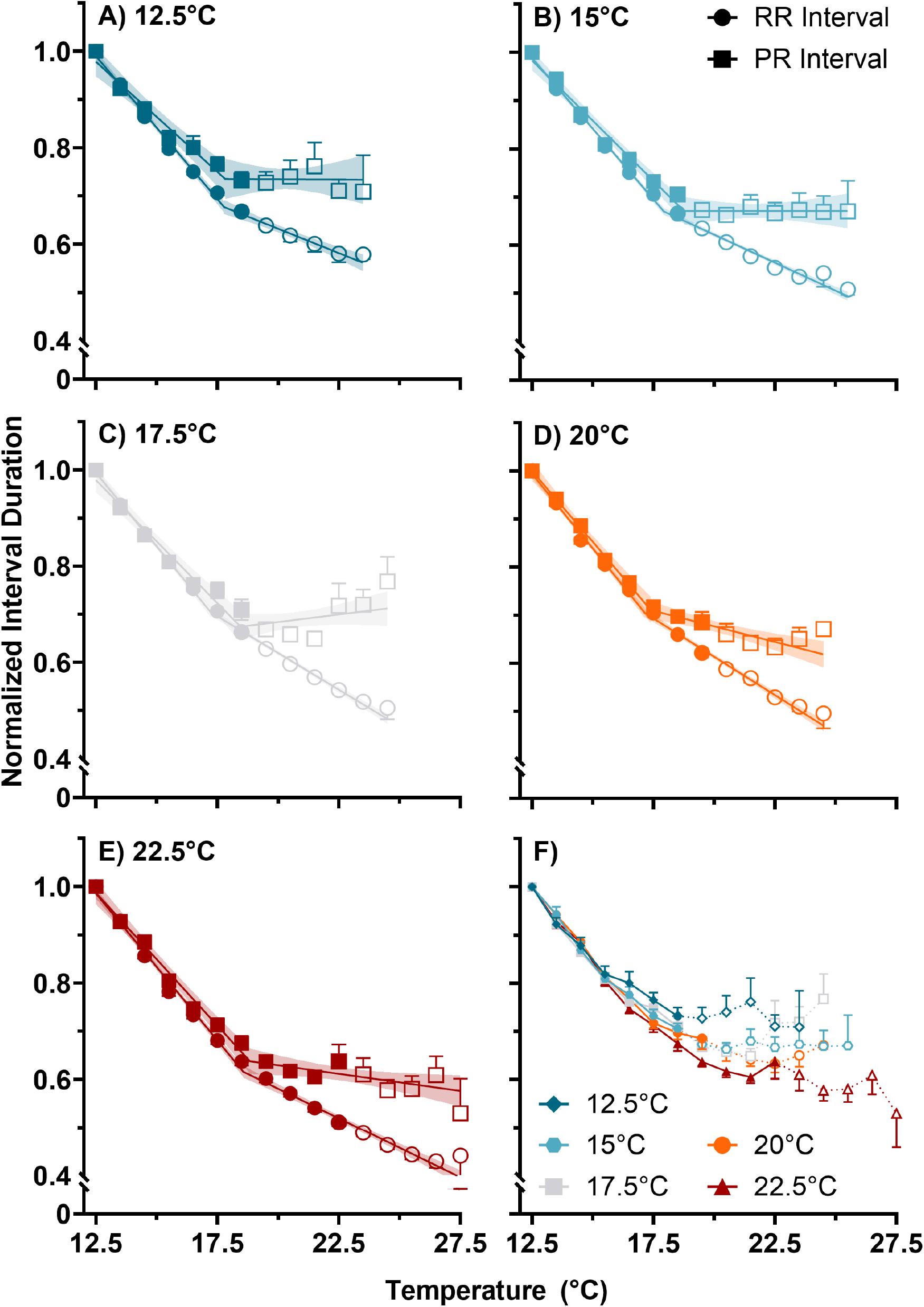
Normalized PR and RR intervals for maximum heart rate during acute warming in juvenile Chinook salmon acclimated to A) 12.5, B) 15, C) 17.5, D) 20, and E) 22.5°C. F) Normalized PR interval for all acclimation temperatures. Values are normalized to the initial value at the onset of warming (12.5°C). Acute warming continued at 10°C hr^-1^ until the onset of arrhythmia. Data are presented as mean ± SE with the respective segmented linear regression (SLR) line and the 95% confidence interval. Open symbols denote point at which individuals had been removed after the onset of arrhythmia. **Alt text:** Graphs depicting the response of the PR and RR intervals with acute warming between five acclimation temperatures with segmented linear regression. Graphs are labeled A to F.

## Discussion

In a rapidly warming north, high-latitude Chinook salmon populations are experiencing greater thermal variation and peak environmental temperatures throughout freshwater life stages (embryos, juveniles, and spawning adults; Von Biela et al., 2020, 2023). We characterized juvenile Yukon River Chinook salmon whole animal and cardiac acute heat tolerance and thermal sensitivity with temperature acclimation. In line with our predictions, Yukon River Chinook salmon showed improvements to whole animal and cardiac heat tolerance with warm acclimation and a marked resetting of heart rate, demonstrating sustained thermal sensitivity and function under acute heat stress. Additionally, performance did not decline with acclimation near and at their peak ecological temperatures (20 and 22.5°C), showing that Yukon River Chinook salmon have a substantial capacity to increase their heat tolerance with acclimation to peak summer water temperatures. This finding supports our second hypothesis that despite their high-latitude distribution, Yukon River Chinook salmon have retained broad thermal tolerance, potentially as a result of high historical thermal variation or a challenging migration.

### Whole animal heat tolerance

Yukon River juvenile Chinook salmon demonstrate a remarkable whole animal heat tolerance and ability to increase their upper thermal limits with warm acclimation, comparable to or exceeding temperate Chinook salmon strains and other salmonids. Yukon River Chinook salmon raised at 12.5°C demonstrated a comparable CT_max_ (28.6°C) to temperate Chinook salmon raised at 11°C (28.0°C; Zillig et al., 2025). At intermediate and warm acclimation temperatures, Yukon River Chinook salmon displayed a CT_max_ ∼1-2°C above that of temperate strains (Zillig et al., 2025). These values are equivalent to the highest CT_max_ values documented in Pacific salmonids (Adams et al., 2022; Butler et al., 2026; Strowbridge et al., 2025; Zillig et al., 2025). Furthermore, the plasticity we observed in CT_max_ was also greater than that typically documented in Pacific salmon. For instance, in temperate Chinook salmon, Zillig et al. (2025) and Butler et al., (2026) found CTmax ARRs of 0.16 between 11 and 16°C and 0.04 between 15 and 18°C respectively, while in rainbow trout, Strowbridge et al. (2025) found an ARR of 0.27 between 12 and 18°C. Here, we found a substantially higher ARR of 0.31 over comparable temperatures (12.5 *vs* 17.5°C). As in other salmonids, plasticity for CT_max_ appeared to plateau above 20°C, suggesting that Yukon River Chinook salmon were approaching their limit for warm acclimation (Butler et al., 2026; Strowbridge et al., 2025; Zillig et al., 2025).

Yukon River Chinook salmon also displayed greater upper thermal tolerance and plasticity than temperate Chinook salmon during exercise. British Columbian (BC) Nicola River Chinook salmon experience similar thermal variation but water temperatures peak ∼4°C higher (∼25°C) than the Yukon River (Butler et al., 2026; AKTEMP, 2026). Both strains had comparable CT_swim_ values when acclimated to 15°C (26.2 and 26.6°C, respectively), but Yukon River Chinook salmon exhibited a CT_swim_ 1-3°C above the temperate strain when acclimated to temperatures above 15°C (Butler et al., 2026). With acclimation to their ecological peak temperature (24°C), CT_swim_ of Nicola River Chinook salmon dropped ∼2°C to 25.5°C, while Yukon River Chinook salmon maintained CT_swim_ at their ecological peak temperature (22.5°C acclimation, CT_swim_ 28.3°C; Butler et al., 2026). As with CT_max_, Yukon River Chinook salmon showed greater plasticity than the temperate strain. Between 15 and 20°C, ARR was 0.33 in Yukon River and 0.22 in Nicola River Chinook salmon (Butler et al., 2026). The ARR for both strains dropped substantially at higher acclimation temperatures, but more so in Nicola River Chinook salmon (Yukon River Chinook: 17.5 *vs* 22.5°C, ARR 0.02; Nicola River Chinook: 18 *vs* 24°C, ARR -0.15; Butler et al., 2026). This suggests that Yukon River Chinook salmon can acclimate to peak temperatures encountered in the wild (21-22°C), and while functional traits, such as swimming performance during acute warming, may not improve with acclimation above the average temperatures they experience (15-18°C), whole organism performance is preserved.

Importantly, while whole animal performance did not decline in our highest acclimation group, growth and feeding rates did (unpublished data), suggesting other physiological costs. Further, the Nicola River Chinook salmon were roughly half the size of the Yukon River Chinook salmon and the warming rate used in this study was substantially faster than that used in Butler et al. (2026). However, even absent a decline in swimming performance, recovery from exhaustive swimming at high temperatures can be impaired in Chinook salmon and other salmonids (Butler et al., 2026; Gilbert & Tierney, 2018). Therefore, future studies of northern Chinook salmon should examine effects of temperature on post-exercise recovery dynamics and survival.

Life history variation, including freshwater residence duration and migration timing and location, may shape salmon thermal tolerance and acclimation capacity. In Chinook salmon, genomic signatures of thermal tolerance clustered primarily by population and life-history type (Metzger et al., 2026). Freshwater residence time, migration distance, slope, and water temperature have also been implicated in shaping genetic responses of salmonids (Metzger et al., 2026; Micheletti et al., 2018). Indeed, Van Wert et al. (2023) and Dressler et al. (2023, 2025) found that fish with warmer, more strenuous migrations exhibited higher CT_max_ and lower mortality, but less plasticity, than fish with cooler, shorter migrations. Cardiac traits, from whole organ to molecular level, also vary with migration life history (Eliason et al., 2011). In particular, populations completing more arduous or warmer migrations can exhibit greater aerobic capacity (Anttila et al., 2014; Farrell et al., 2008; Eliason et al., 2011). Such variation may underly the high heat tolerance of Yukon River Chinook salmon, which spend a full year rearing in freshwater, have the longest migration of any Pacific salmon, and migrate during peak summer water temperatures (∼15-18°C).

### Cardiac heat tolerance and temperature sensitivity

Acclimation to high temperatures commonly improves cardiac heat tolerance under acute warming (e.g. Adams et al., 2022; Anttila, Couturier, et al., 2014; Drost et al., 2016; Gilbert, Adams, et al., 2022; Gilbert et al., 2024; Gilbert & Farrell, 2021; Muñoz et al., 2015; Williams et al., 2025), but the extent of this response varies within and among species. In Quinsam River Chinook salmon, warm acclimation from 8 to 12.4°C increased T_peak_ by ∼2°C but T_arr_ by only 0.3°C, and family reaction norms for T_arr_ ranged from no change to increases of more than 1°C (Muñoz et al., 2015). Similarly, some rainbow trout strains show large increases in cardiac thermal limits with warm acclimation (Adams et al., 2022; Strowbridge et al., 2025), while others do not (Gilbert et al., 2022). Our results fit within this variation. When fed to satiation and acclimated to 12.5°C, Yukon River Chinook salmon had a T_peak_ (22.6°C) within, and a T_arr_ (23.4°C) at the lower end of, the range of family means for Quinsam River Chinook salmon reared at 12.4°C (∼21.9–23.6 and ∼23.5–25.7°C, respectively; Muñoz et al., 2015). Unlike in Quinsam River Chinook salmon, however, T_peak_ and T_arr_ increased in parallel with acclimation in Yukon River Chinook salmon (ARR 0.18–0.19 between 12.5 and 22.5°C), albeit over a warmer range of acclimation temperatures and to a lesser extent than in Atlantic salmon (ARR 0.63 and 0.66 between 12 and 20°C; Anttila et al., 2014).

Together with the plasticity of CT_max_, these responses suggest that the outcomes and mechanisms of plasticity likely vary among thermal tolerance traits, not only among species and populations. Yukon River Chinook salmon had a greater whole animal heat tolerance than more southern Chinook salmon but a comparable or slightly lower cardiac heat tolerance, and the acclimation responses of these traits followed different patterns. The ARR for CT_max_ was greatest between the coolest acclimation temperatures and declined steadily with warming (0.34 to 0.15), while T_peak_ and T_arr_ showed their largest increases between 20 and 22.5°C (ARR 0.62 and 0.63), where plasticity in CT_max_ was lowest. Similarly, Gilbert et al. (2021) found that T_arr_ increased with acclimation far more than CT_max_, such that the two differed markedly after cold acclimation but converged after warm acclimation. At rapid warming rates, CT_max_ may be set largely by neural failure rather than by limits to oxygen delivery and may therefore capture a distinct aspect of plasticity from T_peak_ and T_arr_, which reflect cardiac or cardiorespiratory plasticity specifically. A limited acclimation response in one trait should therefore not be taken as a limited capacity for thermal plasticity overall.

Resetting of *f*_Hmax_ with warm acclimation has been documented to varying degrees in salmonids and other fishes. For example, when *f*_Hmax_ was tested at a common intermediate temperature it was 26% lower in Atlantic salmon acclimated at 20 relative to 12°C and 21–29% lower in Arctic char acclimated to 14°C compared to those acclimated to 2°C (Anttila et al., 2014; Gilbert & Farrell, 2021). Likewise, we found that *f*_Hmax_ at common moderate temperatures decreased progressively as acclimation temperature increased, with 22.5°C acclimated fish showing a reduction in *f*_Hmax_ by ∼15 beats min_-_^1^ at 17.5°C during acute warming compared to 12.5°C acclimated fish. Peak *f*_Hmax_ increased only slightly, so the modest increase in Δ*f*_Hmax_ was driven largely by the reduction in initial *f*_Hmax_ (Fig. 2A-B, Table 2). Thus, our results support the notion that the mechanisms underlying resetting of *f*_Hmax_ are, at least in part, uncoupled from those responsible for increasing peak *f*_Hmax_. Resetting of *f*_Hmax_ is likely dependent on changes in cardiac pacemaker rates, whereas changes in peak *f*_Hmax_ likely depend more on a broader electrical remodeling of the heart that supports excitability at high temperatures. Accordingly, Anttila et al. (2014), Strowbridge et al. (2025), Gilbert & Farrell (2021), and Gilbert et al. (2022) all found substantial increases in peak *f*_Hmax_ in various salmonids following warm acclimation, while like us, Adams et al. (2022) found relatively minor differences in peak *f*_Hmax_ and Δ*f*_Hmax_ with acclimation.

### Electrical activity of the heart

Numerous studies have documented extensive electrical remodeling of the heart with thermal acclimation (e.g. Abramochkin et al., 2019; Aho & Vornanen, 1998; Haverinen & Vornanen, 2009, 2020; Vornanen, 2016, 2021; Vornanen et al., 2002). For instance, a population of Arctic lake trout exhibited decreasing PR, QRS, and QT intervals with acute warming when winter and summer acclimatized (Williams et al., 2025). Here we found that both the PR and QT intervals decreased with acute warming while the QRS interval increased; however, the trends of these changes varied with acclimation temperature.

Decreasing PR interval indicates faster atrioventricular (AV) conduction and corresponds primarily to *I*_Na_ currents, which drive the action potential (AP) upstroke (Vornanen, 2017). Previous work on rainbow trout acclimated to 4°C demonstrated thermal compensation through an increase in *I*_Na_ density to offset the slowing effects of cold on AV conduction (Haverinen and Vornanen, 2004; Vornanen, 2017). In contrast, Yukon River Chinook salmon increased PR interval with acclimation to 12.5°C, suggesting decreasing *I*_Na_ densities with acclimation from 22.5 to 12.5°C. Furthermore, we observed a marked plateau in the PR interval at intermediate temperatures, regardless of acclimation temperature, that may be driven by the high thermal sensitivity of *I*_Na_, ultimately reducing the electrical excitability of the heart (Vornanen, 2017). These results provide support that the functional temperature limits of *I*_Na_ may be an important limiting factor to cardiac function under thermal stress.

Potassium currents (K^+^) are thermally sensitive and play a substantial role in setting the excitability of cardiac myocytes (Vornanen, 2017; Haverinen & Vornanen, 2020). Increases in K^+^ current density (specifically *I*_Kr_) have been shown to drive faster repolarization (shorter QT interval) with cold acclimation (Filatova et al., 2019; Haverinen & Vornanen, 2009; Vornanen, 2016b). Here, Chinook salmon show decreasing QT intervals with acclimation from 22.5 to 12.5°C, indicative of faster repolarization and a potential increase in K ^+^ currents. However, an increase in K^+^ current densities, if not matched by an increase in *I*_Na_, can reduce cardiac excitability (Hassinen et al., 2007; Vornanen, 2017; Vornanen et al., 2002). Therefore, the potential upregulation of K^+^ currents paired with a downregulation of *I*_Na_ may result in a mismatch between the flow of these key ions, limiting excitation (Haverinen & Vornanen, 2020). In humans, increased circulating K^+^ levels have also been linked to prolongation of the QRS interval (Weiss et al., 2017). Here, Chinook salmon shows a prolongation of QRS duration with acclimation from 22.5 to 12.5°C, further indicating a potential upregulation of K^+^ current density and/or circulating K^+^ levels. Combined, these changes indicate marked thermal plasticity in the cardiac electrical activity of Yukon River Chinook salmon.

### Conclusions and ecological implications

Previous work has suggested that heat tolerance may be driven by local adaptation to historical river temperatures which vary by latitude and by migration distance (Zillig et al., 2025, 2023; Muñoz et al., 2015; Anttila et al., 2014; Eliason et al., 2011; Gilbert & Farrell, 2021). We expected that Yukon River salmon could show a limited heat tolerance and capacity for warm acclimation due to their high latitude and historically cool average temperatures. In stark contrast, we found that juvenile Yukon River Chinook salmon possess a strong ability to increase acute heat tolerance with warm acclimation and demonstrate a high acute heat tolerance comparable to or greater than that found in many temperate populations. This thermal performance may instead reflect the extreme thermal variation faced by salmon at high latitudes or have a shared physiological basis with a high-performance phenotype associated with their lengthy and challenging migration. Indeed, others have previously found that Pacific salmon populations faced with more challenging migrations tended to have higher heat tolerance (Eliason et al., 2011).

While this study provides the first assessment of whole animal and cardiac thermal tolerance in juvenile Yukon River Chinook salmon, there are limits to direct ecological relevance of the thermal tolerance values generated. The present study focused on assessing acute heat tolerance using common methodologies with rapid warming rates that are not commonly encountered in nature. Likewise, *f*_Hmax_ assessments reveal factors that may limit oxygen delivery and thus whole organism function during warming, but whole organism function cannot be distilled to a single organ system as other factors contribute (Pörtner et al., 2017). Such values allow rapid characterization of plasticity and comparison across studies, but assessing other functional traits (e.g. growth, food consumption, swimming performance) under simulation of more natural thermal variation and with co-occurring stressors (e.g., disease or hypoxia) would be natural next steps to delineate sustained thermal limits within the acute boundaries for the performance identified here.

Despite these limitations, our work has important conservation implications. First, as shown in other species and populations (e.g. Morley et al., 2019; Zillig et al., 2025), Yukon River Chinook salmon have strong acclimation capacity, so vulnerability assessments and thermal habitat models that do not incorporate that plasticity may overestimate their sensitivity to warming. Second, because acute whole animal heat tolerance was among the highest reported for juvenile Chinook salmon and cardiac thermal limits were comparable to those of temperate strains, high-latitude populations should not be assumed to have reduced heat tolerance. Finally, although Yukon River Chinook salmon are among the northernmost Chinook salmon populations, we found no temperature-dependent mortality at our highest acclimation temperature (22.5°C) over >4 weeks. Thus, Yukon River Chinook salmon may have an upper thermal limit for acclimation above that tested here. This lack of mortality combined with our acute thermal performance data suggests that direct mortality associated with acute heat exposure during freshwater rearing is unlikely to be a major contributor to Yukon River Chinook salmon declines. However, longer exposure to temperatures above 22.5°C, high temperature in combination with co-occurring stressors, and sublethal effects at warm temperatures, such as the reduced feeding and growth we observed at 22.5°C (unpublished data), are all potential limiters to the success of northern juvenile Chinook salmon that require further investigation.

## Acknowledgements

We thank Travis Hyer, the staff at the Ruth Burnett Sport Fish Hatchery, and the Alaska Department of Fish and Game for producing and providing the Chinook salmon used in this study and for information used to improve fish husbandry. We also thank the staff at the University of Alaska Fairbanks Animal Resource Center and Will Samuel for supporting animal husbandry and facilities operations. Research reported was supported by an Institutional Development Award (IDeA) from the National Institute of General Medical Sciences of the National Institutes of Health under grant number P20GM103395 and a Student Project Award from the UAF Undergraduate Research and Scholarly Activity (URSA) program. The content is solely the responsibility of the authors and does not necessarily reflect the official views of the NIH.

