## Supplemental Material for "High latitude, high tolerance: Northern juvenile Chinook salmon exhibit substantial thermal plasticity and acute heat tolerance"

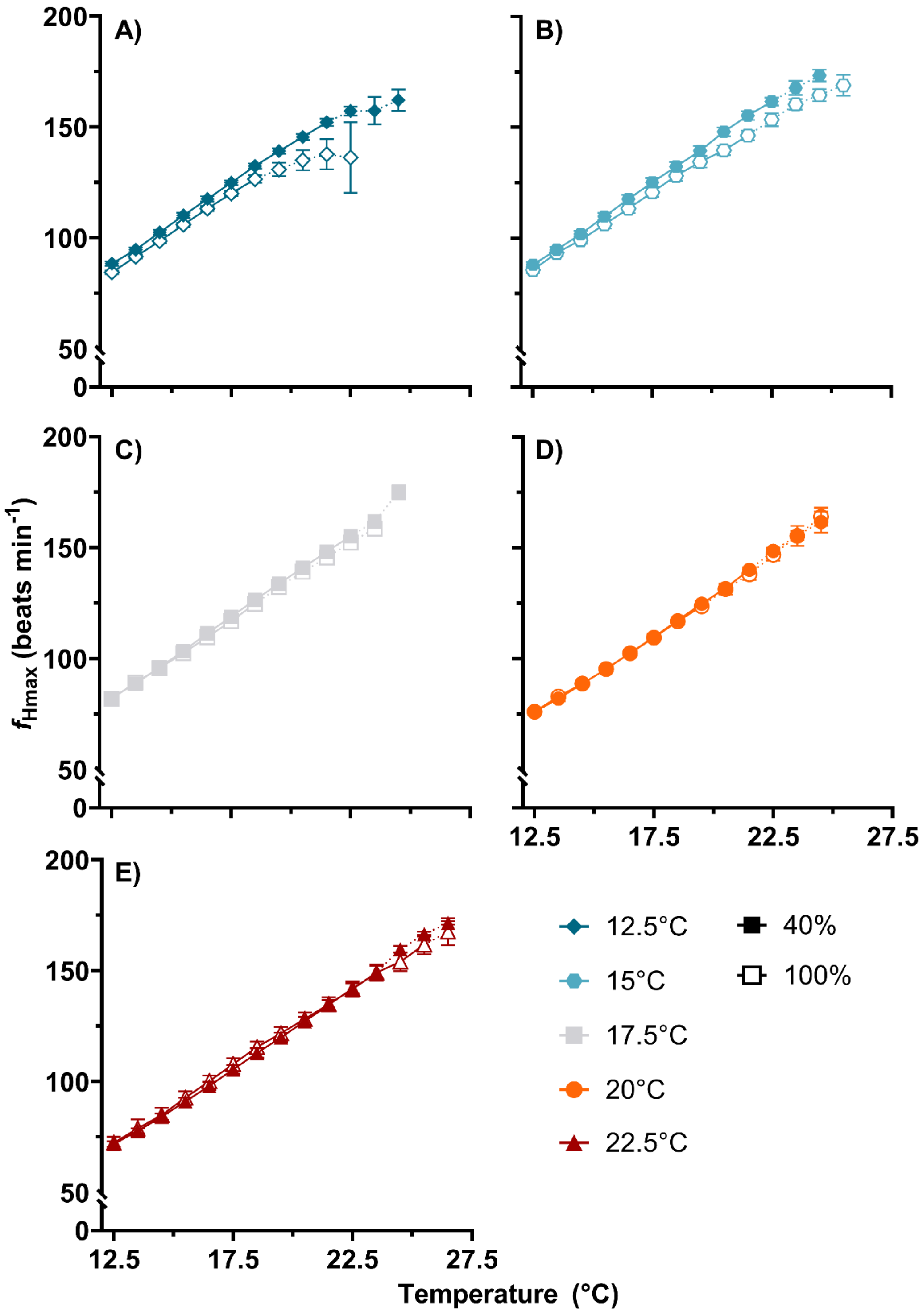


**Figure S1:** Response of maximum heart rate ($f_{Hmax}$) to acute warming for juvenile Yukon River Chinook acclimated to A) 12.5, B) 15, C) 17.5, D) 20, and E) 22.5°C under two feeding treatments of 100% (filled symbols) or 40% of satiation (open symbols). Acute warming began at 12.5°C and continued at 10°C hr$^{-1}$ until the onset of arrhythmia. Data are presented as mean ± SE. The dotted line denotes the point at which two individuals have been removed after the onset of arrhythmia. Model statistics are presented in Table S1.

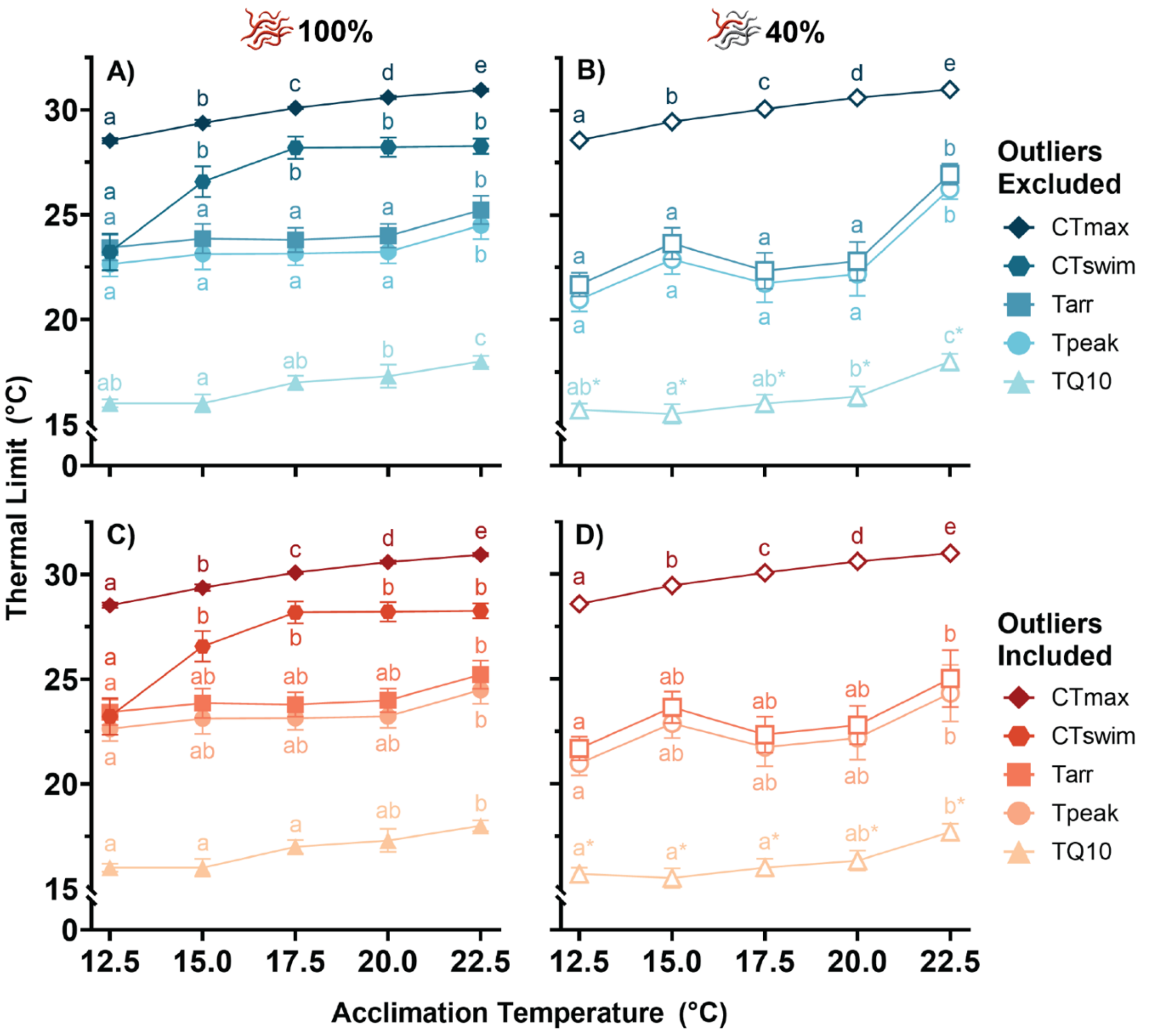


**Figure S2:** Effect of acclimation on transition and critical temperatures for juvenile Yukon River Chinook acclimated to 12.5, 15, 17.5, 20, and 22.5°C and fed to 100% (filled symbols) or 40% of satiation (open symbols). Transition and critical temperatures include critical thermal maxima static ($CT_{max}$) and under exercise ($CT_{swim}$), temperature at peak heart rate ($T_{peak}$), temperature at arrhythmia ($T_{arr}$) and temperature at which incremental $Q_{10}$ dropped and stayed below 2.0 ($T_{Q10}$). Thermal limits are shown for both the quality controlled dataset with two outliers removed (n = 82) due to early indications of poor sample preparation (e.g. variable QRS amplitude) and the complete dataset (n = 84). Data are presented as mean ± SE. Dissimilar letters indicate significant differences between acclimation temperatures and asterisks denote significant differences between feed treatments. Model statistics are presented in Table S3.

**Fixed effects:**

| **Mass** | **Sum of Sq** | **df** | **F** | ***p*** | | |
|---|---|---|---|---|---|---|
| (Intercept) | 2149.25 | 1 | 441.9427 | <0.0001 | | |
| Acclimation | 17.67 | 4 | 0.9085 | 0.4636 | | |
| Feed treatment | 16.37 | 1 | 3.3651 | 0.0706 | | |
| **Length** | **Chisq** | **df** | ***p*** | | | |
| Acclimation | 3.2271 | 4 | 0.5206 | | | |
| Feed treatment | 0.0003 | 1 | 0.9859 | | | |
| **Body condition (K)** | | | | | ***Post-hoc results*** | ***p*** |
| Acclimation | 0.064 | 4 | 0.9995 | | 40% Feed treatment - 100% Feed treatment | 0.0063 |
| Feed treatment | 7.8879 | 1 | 0.005 | | | |

**Model:**

Mass ~ Acclimation + Feed treatment

Length ~ Acclimation + Feed treatment
Body Condition ~ Acclimation + Feed treatment

**Table S1:** ANOVA and post-hoc results for linear and generalized linear models on the body condition of juvenile Yukon River Chinook acclimated to 12.5, 15, 17.5, 20, and 22.5°C and fed to 40 or 100% satiation. Significant effects ($p<0.05$) were calculated using a type III Wald chi-square test and significant fixed effects were subjected to further pairwise comparisons with the Holm adjustment.

**Fixed effects:**

| **peak $f_{Hmax}$** | **Chisq** | **df** | **p** | ***Post-hoc results*** | ***p*** |
|---|---|---|---|---|---|
| Acclimation | 11.9636 | 4 | 0.0176 | Acclimation 12.5°C - Acclimation 15°C | 0.1832 |
| Feed treatment | 4.8482 | 1 | 0.0277 | Acclimation 12.5°C - Acclimation 17.5°C | 1 |
| | | | | Acclimation 12.5°C - Acclimation 20°C | 1 |
| | | | | Acclimation 12.5°C - Acclimation 22.5°C | 0.1219 |
| | | | | Acclimation 15°C - Acclimation 17.5°C | 1 |
| | | | | Acclimation 15°C - Acclimation 20°C | 0.1725 |
| | | | | Acclimation 15°C - Acclimation 22.5°C | 1 |
| | | | | Acclimation 17.5°C - Acclimation 20°C | 1 |

| | | | | | | |
|---|---|---|---|---|---|---|
| | | | | | Acclimation 17.5°C - Acclimation 22.5°C | 0.9216 |
| | | | | | Acclimation 20°C - Acclimation 22.5°C | 0.1124 |
| | | | | | 40% Feed treatment – 100% Feed treatment | 0.0307 |
| **Δ$f_{Hmax}$** | | | | | | |
| Acclimation | 9.4699 | 4 | 0.0504 | | | |
| Feed treatment | 1.8273 | 1 | 0.1764 | | | |
| **$f_{Hmax}$ at 12.5°C** | | | | | ***Post-hoc results*** | ***p*** |
| Acclimation | 173.661 | 4 | <0.0001 | | Acclimation 12.5°C - Acclimation 15°C | 0.6753 |
| Feed treatment | 1.856 | 1 | 0.1731 | | Acclimation 12.5°C - Acclimation 17.5°C | 0.0116 |
| | | | | | Acclimation 12.5°C - Acclimation 20°C | <0.0001 |
| | | | | | Acclimation 12.5°C - Acclimation 22.5°C | <0.0001 |
| | | | | | Acclimation 15°C - Acclimation 17.5°C | 0.0071 |
| | | | | | Acclimation 15°C - Acclimation 20°C | <0.0001 |
| | | | | | Acclimation 15°C - Acclimation 22.5°C | <0.0001 |
| | | | | | Acclimation 17.5°C - Acclimation 20°C | 0.0002 |
| | | | | | Acclimation 17.5°C - Acclimation 22.5°C | <0.0001 |
| | | | | | Acclimation 20°C - Acclimation 22.5°C | 0.0116 |
| **$f_{Hmax}$ at 17.5°C** | **Sum of Sq** | **df** | **F** | **p** | ***Post-hoc results*** | |
| Intercept | 225181 | 1 | 9450.888 | <0.0001 | Acclimation 12.5°C - Acclimation 15°C | 0.6649 |
| Acclimation | 3597 | 4 | 37.739 | <0.0001 | Acclimation 12.5°C - Acclimation 17.5°C | 0.0311 |
| Feed treatment | 73 | 1 | 3.076 | 0.0835 | Acclimation 12.5°C - Acclimation 20°C | <0.0001 |
| | | | | | Acclimation 12.5°C - Acclimation 22.5°C | <0.0001 |
| | | | | | Acclimation 15°C - Acclimation 17.5°C | 0.0155 |

| | | | | | |
|---|---|---|---|---|---|
| | | | | Acclimation 15°C - Acclimation 20°C | <0.0001 |
| | | | | Acclimation 15°C - Acclimation 22.5°C | <0.0001 |
| | | | | Acclimation 17.5°C - Acclimation 20°C | <0.0001 |
| | | | | Acclimation 17.5°C - Acclimation 22.5°C | <0.0001 |
| | | | | Acclimation 20°C - Acclimation 22.5°C | 0.2248 |
| **$f_{Hmax}$ at 22.5°C** | **Chisq** | **df** | **p** | ***Post-hoc results*** | |
| Acclimation | 36.26 | 4 | <0.0001 | Acclimation 12.5°C - Acclimation 15°C | 0.2135 |
| Feed treatment | 5.635 | 1 | 0.0176 | Acclimation 12.5°C - Acclimation 17.5°C | 0.6582 |
| | | | | Acclimation 12.5°C - Acclimation 20°C | 0.6582 |
| | | | | Acclimation 12.5°C - Acclimation 22.5°C | 0.1117 |
| | | | | Acclimation 15°C - Acclimation 17.5°C | 0.5235 |
| | | | | Acclimation 15°C - Acclimation 20°C | 0.0149 |
| | | | | Acclimation 15°C - Acclimation 22.5°C | <0.0001 |
| | | | | Acclimation 17.5°C - Acclimation 20°C | 0.2507 |
| | | | | Acclimation 17.5°C - Acclimation 22.5°C | 0.001 |
| | | | | Acclimation 20°C - Acclimation 22.5°C | 0.2875 |
| | | | | 40% Feed treatment – 100% Feed treatment | 0.0211 |
| **$T_{peak}$** | | | | ***Post-hoc results*** | |
| Acclimation | 30.5029 | 4 | <0.0001 | Acclimation 12.5°C - Acclimation 15°C | 0.3645 |
| Feed treatment | 1.8463 | 1 | 0.1742 | Acclimation 12.5°C - Acclimation 17.5°C | 1 |
| | | | | Acclimation 12.5°C - Acclimation 20°C | 0.6403 |

| | | | | | | |
|---|---|---|---|---|---|---|
| | | | | | Acclimation 12.5°C - Acclimation 22.5°C | <0.0001 |
| | | | | | Acclimation 15°C - Acclimation 17.5°C | 1 |
| | | | | | Acclimation 15°C - Acclimation 20°C | 1 |
| | | | | | Acclimation 15°C - Acclimation 22.5°C | 0.0132 |
| | | | | | Acclimation 17.5°C - Acclimation 20°C | 1 |
| | | | | | Acclimation 17.5°C - Acclimation 22.5°C | 0.0012 |
| | | | | | Acclimation 20°C - Acclimation 22.5°C | 0.0052 |
| **$T_{arr}$** | **Sum of Sq** | **df** | **F** | **p** | ***Post-hoc results*** | |
| Intercept | 7537.4 | 1 | 1943.3626 | <0.0001 | Acclimation 12.5°C - Acclimation 15°C | 0.3901 |
| Acclimation | 124.7 | 4 | 8.0361 | <0.0001 | Acclimation 12.5°C - Acclimation 17.5°C | 1 |
| Feed treatment | 7.1 | 1 | 1.8352 | 0.1795 | Acclimation 12.5°C - Acclimation 20°C | 0.7713 |
| | | | | | Acclimation 12.5°C - Acclimation 22.5°C | <0.0001 |
| | | | | | Acclimation 15°C - Acclimation 17.5°C | 1 |
| | | | | | Acclimation 15°C - Acclimation 20°C | 1 |
| | | | | | Acclimation 15°C - Acclimation 22.5°C | 0.0089 |
| | | | | | Acclimation 17.5°C - Acclimation 20°C | 1 |
| | | | | | Acclimation 17.5°C - Acclimation 22.5°C | 0.0004 |
| | | | | | Acclimation 20°C - Acclimation 22.5°C | 0.0028 |
| **$T_{Q10}$** | | | | | ***Post-hoc results*** | |
| Intercept | 3720.9 | 1 | 2915.6489 | <0.0001 | Acclimation 12.5°C - Acclimation 15°C | 0.771 |
| Acclimation | 53.9 | 4 | 10.5565 | <0.0001 | Acclimation 12.5°C - Acclimation 17.5°C | 0.3162 |

| | | | | | | |
|---|---|---|---|---|---|---|
| Feed treatment | 5.9 | 1 | 4.6595 | 0.034 | Acclimation 12.5°C - Acclimation 20°C | 0.0599 |
| | | | | | Acclimation 12.5°C - Acclimation 22.5°C | <0.0001 |
| | | | | | Acclimation 15°C - Acclimation 17.5°C | 0.257 |
| | | | | | Acclimation 15°C - Acclimation 20°C | 0.0427 |
| | | | | | Acclimation 15°C - Acclimation 22.5°C | <0.0001 |
| | | | | | Acclimation 17.5°C - Acclimation 20°C | 0.7182 |
| | | | | | Acclimation 17.5°C - Acclimation 22.5°C | 0.0027 |
| | | | | | Acclimation 20°C - Acclimation 22.5°C | 0.0427 |
| | | | | | 40% Feed treatment – 100% Feed treatment | 0.034 |
| **$CT_{max}$** | **Chisq** | **df** | **p** | | ***Post-hoc results*** | |
| Acclimation | 933.27 | 4 | <0.0001 | | Acclimation 12.5°C - Acclimation 15°C | <0.0001 |
| Feed treatment | 0.29 | 1 | 0.5911 | | Acclimation 12.5°C - Acclimation 17.5°C | <0.0001 |
| | | | | | Acclimation 12.5°C - Acclimation 20°C | <0.0001 |
| | | | | | Acclimation 12.5°C - Acclimation 22.5°C | <0.0001 |
| | | | | | Acclimation 15°C - Acclimation 17.5°C | <0.0001 |
| | | | | | Acclimation 15°C - Acclimation 20°C | <0.0001 |
| | | | | | Acclimation 15°C - Acclimation 22.5°C | <0.0001 |
| | | | | | Acclimation 17.5°C - Acclimation 20°C | <0.0001 |
| | | | | | Acclimation 17.5°C - Acclimation 22.5°C | <0.0001 |
| | | | | | Acclimation 20°C - Acclimation 22.5°C | 0.0001 |
| **$CT_{swim}$** | | | | | ***Post-hoc results*** | |

| | | | | | |
|---|---|---|---|---|---|
| Acclimation | 46.149 | 4 | <0.0001 | Acclimation 12.5°C - Acclimation 15°C | 0.0023 |

**Model:**

Peak $f_{Hmax}$ ~ Acclimation + Feed treatment + 1

$\Delta f_{Hmax}$ ~ Acclimation + Feed treatment

BPM ~ Acclimation + Feed treatment

BPM ~ Acclimation + Feed treatment

BPM ~ Acclimation + Feed treatment + 1

$T_{peak}$ ~ Acclimation + Feed treatment

$T_{arr}$ ~ Acclimation + Feed treatment

$T_{Q10}$ ~ Acclimation + Feed treatment + 1

$CT_{max}$ ~ Acclimation + Feed treatment

$CT_{swim}$ ~ Acclimation + 1

**Table S2:** ANOVA and post-hoc results for linear and generalized linear models on characteristics of cardiac and whole animal function in maximally stimulated juvenile Yukon River Chinook exposed to acute warming. Chinook were acclimated to 12.5, 15, 17.5, 20, and 22.5°C and fed to 40 or 100% satiation. Significant effects (p<0.05) were calculated using a type III Wald chi-square test and significant fixed effects were subjected to further pairwise comparisons with the Holm adjustment.

**Fixed effects:**

| $f_{Hmax}$ | **Coefficient** | **SE** | ***t*** | ***p*** |
|---|---|---|---|---|
| Intercept | 3.8033 | 0.021 | 180.939 | <0.0001 |
| Acclimation 15°C | 0.0037 | 0.0276 | 0.136 | 0.8919 |
| Acclimation 17.5°C | -0.1115 | 0.0277 | -4.022 | <0.0001 |
| Acclimation 20°C | -0.2333 | 0.0278 | -8.385 | <0.0001 |
| Acclimation 22.5°C | -0.2435 | 0.0273 | -8.936 | <0.0001 |
| Acute temperature | 0.0552 | 0.0009 | 61.217 | <0.0001 |
| 100% Feed treatment | -0.0157 | 0.0173 | -0.907 | 0.3645 |
| Acclimation 15°C:Acute temperature | 0.0002 | 0.0011 | 0.225 | 0.8217 |
| Acclimation 17.5°C:Acute temperature | 0.0045 | 0.0011 | 3.98 | <0.0001 |
| Acclimation 20°C:Acute temperature | 0.0075 | 0.0011 | 6.675 | <0.0001 |
| Acclimation 22.5°C:Acute temperature | 0.0058 | 0.001 | 5.728 | <0.0001 |
| Acute temperature:100% Feed treatment | 0.0019 | 0.0006 | 2.887 | 0.0039 |
| ***Post-hoc results*** | | | ***p*** | |
| **Acute temperature** | | **12.5°C** | **17.5°C** | **22.5°C** |
| Acclimation 12.5°C - Acclimation 15°C | | 0.7395 | 0.6846 | 1 |
| Acclimation 12.5°C - Acclimation 17.5°C | | 0.0211 | 0.1984 | 1 |
| Acclimation 12.5°C - Acclimation 20°C | | <0.0001 | <0.0001 | 0.0114 |
| Acclimation 12.5°C - Acclimation 22.5°C | | <0.0001 | <0.0001 | <0.0001 |
| Acclimation 15°C - Acclimation 17.5°C | | 0.0149 | 0.198 | 1 |

| | | | | |
|---|---|---|---|---|
| Acclimation 15°C - Acclimation 20°C | | <0.0001 | <0.0001 | 0.0039 |
| Acclimation 15°C - Acclimation 22.5°C | | <0.0001 | <0.0001 | <0.0001 |
| Acclimation 17.5°C - Acclimation 20°C | | 0.0004 | 0.0039 | 0.0528 |
| Acclimation 17.5°C - Acclimation 22.5°C | | <0.0001 | <0.0001 | <0.0001 |
| Acclimation 20°C - Acclimation 22.5°C | | 0.3059 | 0.198 | 0.1051 |
| **Incremental Q10** | **Coefficient** | **SE** | ***t*** | ***p*** |
| Intercept | 1.2571 | 0.0236 | 53.167 | <0.0001 |
| Acclimation 15°C | 0.006 | 0.0214 | 0.281 | 0.7785 |
| Acclimation 17.5°C | 0.0402 | 0.0215 | 1.872 | 0.0613 |
| Acclimation 20°C | 0.0635 | 0.0216 | 2.942 | 0.0033 |
| Acclimation 22.5°C | 0.1016 | 0.021 | 4.836 | <0.0001 |
| Acute temperature | -0.0385 | 0.001 | -37.362 | <0.0001 |
| 100% Feed treatment | 0.0085 | 0.0138 | 0.614 | 0.5393 |
| ***Post-hoc results*** | | | ***p*** | |
| **Acute temperature** | | **12.5°C** | **17.5°C** | **22.5°C** |
| Acclimation 12.5°C - Acclimation 15°C | | 0.7665 | 0.7665 | 0.7665 |
| Acclimation 12.5°C - Acclimation 17.5°C | | 0.3023 | 0.3023 | 0.3023 |
| Acclimation 12.5°C - Acclimation 20°C | | 0.0214 | 0.0214 | 0.0214 |
| Acclimation 12.5°C - Acclimation 22.5°C | | <0.0001 | <0.0001 | <0.0001 |
| Acclimation 15°C - Acclimation 17.5°C | | 0.4149 | 0.4149 | 0.4149 |
| Acclimation 15°C - Acclimation 20°C | | 0.0499 | 0.0499 | 0.0499 |
| Acclimation 15°C - Acclimation 22.5°C | | 0.0002 | 0.0002 | 0.0002 |
| Acclimation 17.5°C - Acclimation 20°C | | 0.5471 | 0.5471 | 0.5471 |
| Acclimation 17.5°C - Acclimation 22.5°C | | 0.0463 | 0.0463 | 0.0463 |
| Acclimation 20°C - Acclimation 22.5°C | | 0.4149 | 0.4149 | 0.4149 |

| **Model:** | **Conditional $R^2$** | **Marginal $R^2$** |
|---|---|---|
| BPM ~ Acclimation + Acute temperature + Feed treatment + Acclimation x Acute temperature + Acute temperature x Feed treatment + (1 \| Fish ID) | 0.997 | 0.1 |
| Q10 ~ Acclimation + Acute temperature + Feed treatment + (1 \| Fish ID) | 0.912 | 0.11 |

**Table S3:** Generalized linear mixed effect models and post-hoc results characterizing the response of maximum heart rate ($f_{Hmax}$) and incremental $Q_{10}$ to acute warming in juvenile Yukon River Chinook acclimated to 12.5, 15, 17.5, 20, and 22.5°C and fed to 40 or 100% satiation. $f_{Hmax}$ was modelled from 12.5°C and incremental from 13.5°C with fish ID included as a random effect. Significant fixed effects (p<0.05) were subjected to further pairwise comparisons with the Holm adjustment.

**Fixed effects:**

| **PR Interval** | **Coefficient** | **SE** | ***t*** | ***p*** |
|---|---|---|---|---|
| Intercept | -1.6708 | 0.0487 | -34.274 | <0.0001 |

| | | | | |
|---|---|---|---|---|
| Acclimation 15°C | 0.0019 | 0.0655 | 0.029 | 0.9765 |
| Acclimation 17.5°C | -0.0133 | 0.0593 | -0.224 | 0.8228 |
| Acclimation 20°C | 0.0968 | 0.0701 | 1.382 | 0.1669 |
| Acclimation 22.5°C | 0.1673 | 0.0795 | 2.105 | 0.0353 |
| Acute temperature | -0.0302 | 0.0023 | -13.419 | <0.0001 |
| 100% Feed treatment | -0.0455 | 0.0291 | -1.561 | 0.1184 |
| Acclimation 15°C:Acute temperature | -0.0029 | 0.003 | -0.976 | 0.3288 |
| Acclimation 17.5°C:Acute temperature | -0.0031 | 0.0027 | -1.16 | 0.2462 |
| Acclimation 20°C:Acute temperature | -0.0085 | 0.0032 | -2.636 | 0.0084 |
| Acclimation 22.5°C:Acute temperature | -0.0138 | 0.0032 | -4.265 | <0.0001 |
| ***Post-hoc results*** | | | ***p*** | |
| **Acute temperature** | | **12.5°C** | **17.5°C** | **22.5°C** |
| Acclimation 12.5°C - Acclimation 15°C | | 1 | 1 | 1 |
| Acclimation 12.5°C - Acclimation 17.5°C | | 1 | 0.7247 | 0.3443 |
| Acclimation 12.5°C - Acclimation 20°C | | 1 | 1 | 0.3443 |
| Acclimation 12.5°C - Acclimation 22.5°C | | 1 | 1 | 0.1258 |
| Acclimation 15°C - Acclimation 17.5°C | | 1 | 1 | 1 |
| Acclimation 15°C - Acclimation 20°C | | 1 | 1 | 1 |
| Acclimation 15°C - Acclimation 22.5°C | | 1 | 1 | 1 |
| Acclimation 17.5°C - Acclimation 20°C | | 1 | 1 | 1 |
| Acclimation 17.5°C - Acclimation 22.5°C | | 1 | 1 | 1 |
| Acclimation 20°C - Acclimation 22.5°C | | 1 | 1 | 1 |
| **QRS Interval** | **Coefficient** | **SE** | ***t*** | ***p*** |
| Intercept | -2.9828 | 0.0245 | -121.532 | <0.0001 |
| Acclimation 15°C | 0.1123 | 0.0338 | 3.327 | 0.0009 |
| Acclimation 17.5°C | 0.0346 | 0.0305 | 1.135 | 0.2563 |
| Acclimation 20°C | 0.0065 | 0.0361 | 0.181 | 0.8563 |
| Acclimation 22.5°C | 0.0535 | 0.0392 | 1.365 | 0.1721 |
| Acute temperature | 0.0066 | 0.0012 | 5.296 | <0.0001 |
| Acclimation 15°C:Acute temperature | -0.0061 | 0.0016 | -3.714 | 0.0002 |
| Acclimation 17.5°C:Acute temperature | -0.0029 | 0.0015 | -1.933 | 0.0533 |
| Acclimation 20°C:Acute temperature | -0.0022 | 0.0018 | -1.222 | 0.2216 |
| Acclimation 22.5°C:Acute temperature | -0.0042 | 0.0018 | -2.346 | 0.019 |
| ***Post-hoc results*** | | | ***p*** | |
| **Acute temperature** | | **12.5°C** | **17.5°C** | **22.5°C** |
| Acclimation 12.5°C - Acclimation 15°C | | 0.5587 | 1 | 1 |
| Acclimation 12.5°C - Acclimation 17.5°C | | 1 | 1 | 0.857 |
| Acclimation 12.5°C - Acclimation 20°C | | 1 | 0.7949 | 0.406 |
| Acclimation 12.5°C - Acclimation 22.5°C | | 1 | 1 | 0.857 |
| Acclimation 15°C - Acclimation 17.5°C | | 0.4308 | 1 | 1 |

| | | | | |
|---|---|---|---|---|
| Acclimation 15°C - Acclimation 20°C | | 0.0732 | 0.5642 | 1 |
| Acclimation 15°C - Acclimation 22.5°C | | 1 | 1 | 1 |
| Acclimation 17.5°C - Acclimation 20°C | | 1 | 1 | 1 |
| Acclimation 17.5°C - Acclimation 22.5°C | | 1 | 1 | 1 |
| Acclimation 20°C - Acclimation 22.5°C | | 1 | 1 | 1 |
| **QT Interval** | **Coefficient** | **SE** | ***t*** | ***p*** |
| Intercept | -0.5529 | 0.0931 | -5.941 | <0.0001 |
| Acclimation 15°C | 0.0384 | 0.1035 | 0.371 | 0.711 |
| Acclimation 17.5°C | 0.1401 | 0.0948 | 1.478 | 0.139 |
| Acclimation 20°C | 0.1576 | 0.1071 | 1.471 | 0.141 |
| Acclimation 22.5°C | 0.1482 | 0.128 | 1.158 | 0.247 |
| Acute temperature | -0.058 | 0.0029 | -19.875 | <0.0001 |

| **Model:** | **Conditional $R^2$** | **Marginal $R^2$** |
|---|---|---|
| PR Interval ~ Acclimation + Acute temperature + Feed treatment + Acclimation x Acute temperature + (1 \| Fish ID) | 0.055 | 0.966 |
| QRS Interval ~ Acclimation + Acute temperature + Acclimation x Acute temperature + (1 \| Fish ID) | 0.99 | 0.001 |
| QT Interval ~ Acclimation + Acute temperature + Feed treatment + (1 \| Fish ID) | 0.669 | 0.1 |

**Table S4:** Generalized linear mixed effect models and post-hoc results characterizing the response of PR, QRS, and QT interval duration to acute warming in maximally stimulated juvenile Yukon River Chinook acclimated to 12.5, 15, 17.5, 20, and 22.5°C and fed to 40 or 100% satiation. Intervals were modelled from 12.5°C with Fish ID included as a random effect. Significant fixed effects (p<0.05) were subjected to further pairwise comparisons with the Holm adjustment.

| | | **Acclimation Temperature (°C)** | | | | |
|---|---|---|---|---|---|---|
| | | **12.5°C** | **15°C** | **17.5°C** | **20°C** | **22.5°C** |
| **$T_{arr}$** | | $22.5 \pm 0.5^{a}$ | $23.8 \pm 0.5^{a}$ | $23.1 \pm 0.5^{a}$ | $23.5 \pm 0.5^{a}$ | $25.1 \pm 0.8^{b}$ |
| | **40%** | 21.7 ± 0.6 | 23.7 ± 0.8 | 22.3 ± 0.9 | 22.8 ± 0.9 | 25.0 ± 1.4 |
| | **100%** | 23.4 ± 0.6 | 23.9 ± 0.7 | 23.8 ± 0.6 | 24.0 ± 0.6 | 25.2 ± 0.7 |
| **$T_{peak}$** | | $21.7 \pm 0.4^{a}$ | $23.0 \pm 0.5^{a}$ | $22.5 \pm 0.5^{a}$ | $22.8 \pm 0.5^{a}$ | $24.4 \pm 0.8^{b}$ |
| | **40%** | 21.0 ± 0.6 | 22.9 ± 0.7 | 21.8 ± 0.9 | 22.2 ± 1.0 | 24.3 ± 1.4 |
| | **100%** | 22.6 ± 0.6 | 23.1 ± 0.7 | 23.2 ± 0.6 | 23.2 ± 0.6 | 24.5 ± 0.7 |
| **$T_{Q10}$** | | $15.8 \pm 0.2^{ab}$ | $15.8 \pm 0.3^{a}$ | $16.5 \pm 0.3^{ab}$ | $16.9 \pm 0.4^{b}$ | $17.8 \pm 0.2^{c}$ |
| | **40%** | 15.7 ± 0.3* | 15.5 ± 0.5* | 16.0 ± 0.4* | 16.3 ± 0.5* | 17.7 ± 0.4* |
| | **100%** | 16.0 ± 0.2 | 16.0 ± 0.4 | 17.0 ± 0.3 | 17.3 ± 0.6 | 18.0 ± 0.3 |

**Table S5:** Temperature at arrhythmia, peak $f_{Hmax}$, and when incremental $Q_{10}$ falls and remains below 2.0 for juvenile Chinook salmon fed to 40 or 100% satiation and acclimated to 12.5, 15, 17.5, 20, and 22.5°C. Acute warming began at 12.5°C and continued until the onset of arrhythmia. Data are presented as mean ± SE. Model statistics are presented in Table S3.
